# Population connectivity predicts vulnerability to white-nose syndrome in the Chilean myotis (*Myotis chiloensis*) - a genomics approach

**DOI:** 10.1101/2020.04.07.029256

**Authors:** Thomas M. Lilley, Tiina M. Sävilammi TM, Gonzalo Ossa, Anna S Blomberg, Anti Vasemägi, Veronica Yung, David L.J. Vendrami, Joseph S Johnson

## Abstract

Despite its peculiar distribution, the biology of the southernmost bat species in the world, the Chilean myotis (*Myotis chiloensis*), has garnered little attention so far. The species has a north-south distribution of c. 2800 km, mostly on the eastern side of the Andes mountain range. Use of extended torpor occurs in the southernmost portion of the range, putting the species at risk of bat white-nose syndrome (WNS), a fungal disease responsible for massive population declines in North American bats. Here, we examined how geographic distance and topology would be reflected in the population structure of *M. chiloensis* along the majority of its range using a double digestion RAD-tag method. We sampled 66 individuals across the species range and discovered pronounced isolation-by-distance. Furthermore, and surprisingly, we found higher degrees of heterozygosity in the southernmost populations compared to the north. A coalescence analysis revealed that our populations may still not have reached secondary contact after the Last Glacial Maximum. As for the potential spread of pathogens, such as the fungus causing WNS, connectivity among populations was noticeably low, especially between the southern hibernatory populations in the Magallanes and Tierra del Fuego, and more northerly populations. This suggests the probability of geographic spread of the disease from the north through bat-to-bat contact to susceptible populations is low. The study presents a rare case of defined population structure in a bat species and warrants further research on the underlying factors contributing to this.

## INTRODUCTION

Transmission of infectious diseases has garnered attention as one of the greatest risks to human, agriculture and wildlife health over the last decade (Cangelosi *et al*. 2004; Semenza and Menne 2009). Previous research demonstrates that the emergence of previously unknown diseases often results from a change in the ecology of the host, pathogen, and/or their environment (Scholthof 2007). An example of this is white-nose syndrome (hereafter WNS), an epizootic disease that emerged in North America in 2006 (Blehert *et al*. 2009). The disease is caused by the fungus, *Pseudogymnoascus destructans*, which infects insectivorous bats during the hibernation period at latitudes where prey are not widely available during winter (Lorch *et al*. 2011). Populations of highly susceptible species, especially from the genus *Myotis*, have declined by >90% in areas affected by WNS (Frick *et al*. 2010). The opportunistic pathogen can utilize alternative carbon sources (Raudabaugh and Miller 2013) and can persist in the cold, humid environment within hibernacula in the absence of bat hosts (Lorch *et al*. 2013; Hoyt *et al*. 2014).

*P. destructans* is native to Eurasia, where it has a large geographic range, with transmission to North America likely facilitated by humans (Warnecke *et al*. 2012; Leopardi *et al*. 2015). In North America, bats suffering from WNS were first detected in the state of New York during the winter of 2006–2007 (Frick *et al*. 2010). The fungus has since spread across North America, with records of prevalence in 33 U.S. states and 7 Canadian provinces. So far, *P. destructans* has been detected on 17 species of bats, with more species likely to follow. While human-assisted transmission of *P. destructans* likely has contributed to this spread, the ecology and behaviour of cave-hibernating bats in North America also makes them efficient vectors over large geographic areas (Wilder *et al*. 2015). Because WNS affects bats during extended bouts of torpor, at low temperatures where it is able to grow and infect the host, there has been speculation over how far into the southern North America the disease will spread (Verant *et al*. 2012; Meierhofer *et al*. 2019). Although bats inhabiting lower latitudes may suffer less from WNS, *P. destructans* conidia may be able to survive on the body of bats for several months, even at temperatures up to 37 °C (Campbell *et al*. 2019). This could facilitate the movement of WNS across Mesoamerica and the tropics, to arrive to high southern latitudes where bats may be susceptible (Holz *et al*. 2019; Turbill and Welbergen 2019).

Of species known to harbour the WNS fungus, *Tadarida brasiliensis* is of particular interest. As a long-range migratory species, with movements spanning thousands of kilometres (Cockrum 1969; Glass 1982), *T. brasiliensis* may be an important vector for spreading *P. destructans* into the southern hemisphere (Ommundsen *et al*. 2017; McCracken *et al*. 2018). Ecological niche models predict suitable habitat for the proliferation of *P. destructans* in South America, highlighting the need to understand vectors such as *T*. *brasiliensis* as well as human transmission (Escobar *et al*. 2014). However, once *P. destructans* arrives in South America, its spread will not necessarily resemble that seen in North America, as it is likely to be influenced by differing geology and species ecology.

The Chilean myotis (*Myotis chiloensis* [Waterhouse, 1840]) is the most Southerly distributed species of bat in the world, together with the southern big-eared brown bat (*Histiotus magellanicus*, Koopman 1967; Gardner 2007). *Myotis chiloensis* has a vast north-south distribution that includes forested areas on both sides of the Andes from the northern shore of Navarino Island to the southern border of the Atacama desert in Chile (Ossa and Rodriguez-San Pedro 2015). The range of *M. chiloensis* overlaps with the distribution of *T. brasiliensis* from the north, where *M. chiloensis* is not believed to hibernate, to 45°S latitude, where *M. chiloensis* possibly hibernate and may therefore be susceptible to WNS (Bozinovic *et al*. 1985). However, there is no information available on the population structure of *M. chiloensis*, precluding an understanding of how *P. destructans* could be transported from the northern edge of its range to more southern, and vulnerable, populations. The connectedness of individuals across the range will determine the speed and intensity of potential spread. Population structuring in bats is often relatively low because of their efficient mode of dispersal, flight (Laine *et al*. 2013). An ability to disperse more efficiently results in decreased population differentiation (Bohonak 1999) to the extent that some bat species are panmictic across their range (Burland and Wilmer 2001; Laine *et al*. 2013). Such high dispersal would likely result in rapid spread of *P. destructans*. However, bats in the genus *Myotis* show instances of pronounced population structure that may hinder spread of the fungus. For instance, the Gibraltar Strait, which separates the Iberian Peninsula from the Maghreb in Morocco by a minimum gap of 14 km of the open sea, represents a barrier for gene flow for *M. myotis* (Castella *et al*. 2000). Chile is littered with such potential barriers to gene flow, such as the Atacama Desert, glaciers, ice fields, the Andes Mountains, and the Magellan Strait, which in turn can hinder the potential spread of *P. destructans*. Furthermore, populations may still be affected by the Last Glacial Maximum, which covered a large part of Patagonia under ice until c. 10000 years bp (Sérsic *et al*. 2011; Mansilla *et al*. 2018).

With tourism in southern Chile expected to increase (e.g. http://www.conaf.cl/parques-nacionales/visitanos/estadisticas-de-visitacion/), and migratory species such as *T. brasiliensis* capable of carrying spores across large distances, there is a serious need to better understand the population structure of Patagonian bat species before WNS spreads to the region. The lack of knowledge on the extent of migration and mixing, and life history traits in general, means that research in this area is now urgent and essential (Ossa and Rodriguez-San Pedro 2015; Ossa 2016; Ossa *et al*. 2019). Studying population ecology through molecular genetic methods allows for the identification of more accurate population boundaries, which is important when assessing conservation in response to threats of disease and dramatic declines in population size (Moritz 1994). This study will therefore aim to describe population structure and isolation-by-distance in *M. chiloensis* across the range of the species. By studying *M. chiloensis* along 2400 km of latitudinal gradient using genome-wide SNP markers, we aimed to test if geography and the Last Glacial Maximum influence genetic isolation patterns.

## MATERIALS AND METHODS

### Sample collection and DNA extraction

To describe population genetic structure in *M. chiloensis*, we obtained wing tissue samples of 66 bats from two sources. A portion were obtained from live bats captured in the field during November and December 2017 (i.e. austral spring) from two localities: Chicauma, Metropolitana region (33 °S 70 °W) and Karukinka Reserve, Tierra del Fuego (64 °S 78 °W) respectively (Capture permit: 4924_2017, Figure 1 A, Table S1). We used disposable biopsy punches (5 mmm, MLT3335, Miltex Instrument Co, Plainsboro, New Jersey) to collect tissue samples from the plagiopatagium of captured, live bats. The sampled bats were released at the capture site. Additional samples were obtained from dead bats submitted to the Public Health Institute of Chile for rabies testing (Figure 1 A, Table S1). Submitted bats included latitude and longitude locations of origin. Tissue samples from the bats submitted for rabies testing were obtained from the plagiopatium using sterile scalpels. To determine if *P. destructans* had already spread to Chile, we swabbed the nose and wings of all bats in the field and at the Public Health Institute of Chile bat with a sterile polyester swab (Puritan 25-806 1PD, Guildford ME, USA) and stored at -20 °C until analysis.

**Figure 1.**
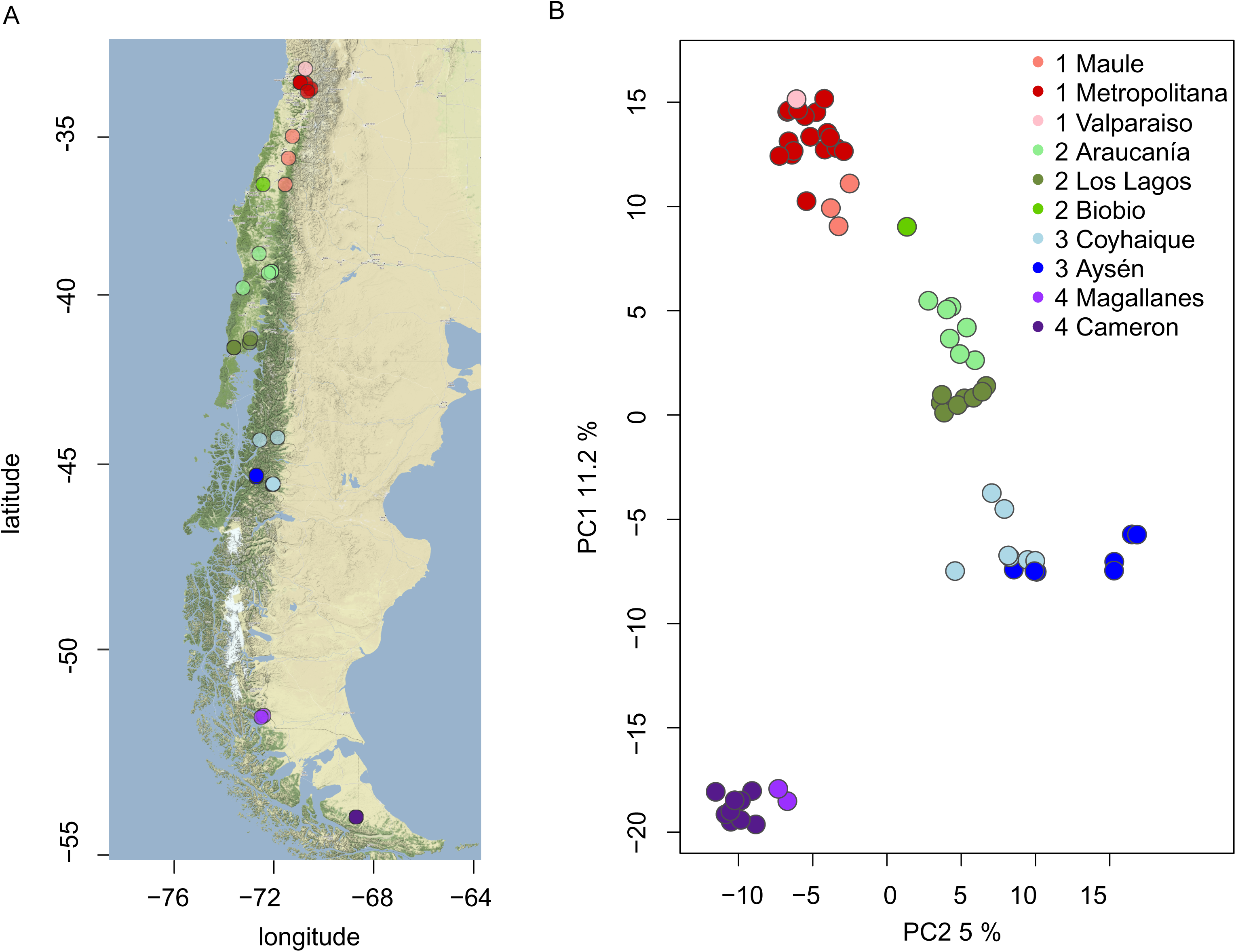
Sampling locations and groupings for genetic sampling of *Myotis chiloensis* in Chile (A). The two most important principal components calculated from allele frequencies explain 15.8% of the total nucleotide variation (B).

We divided samples into four groups according to their geographic origin, with sub-locations within each group to assist in further analyses. Sampling locations are presented in Figure 1 A and details of populations and samples are provided in Table 1 and Table S1. Tissue samples were stored in 1.5 ml tubes with 95% EtOH and stored at -20 ° until further analysis. Fungal spore samples were stored in 1.5 ml tubes and stored at -20 °C until further analysis. We extracted DNA from tissue samples using QIAmp DNA Mini Kits (Qiagen, Hilden Germany) and stored DNA at -80 °. DNA from fungal swabs was extracted using QIamp DNA Micro Kits (Qiagen, Hilden Germany).

**Table 1.**
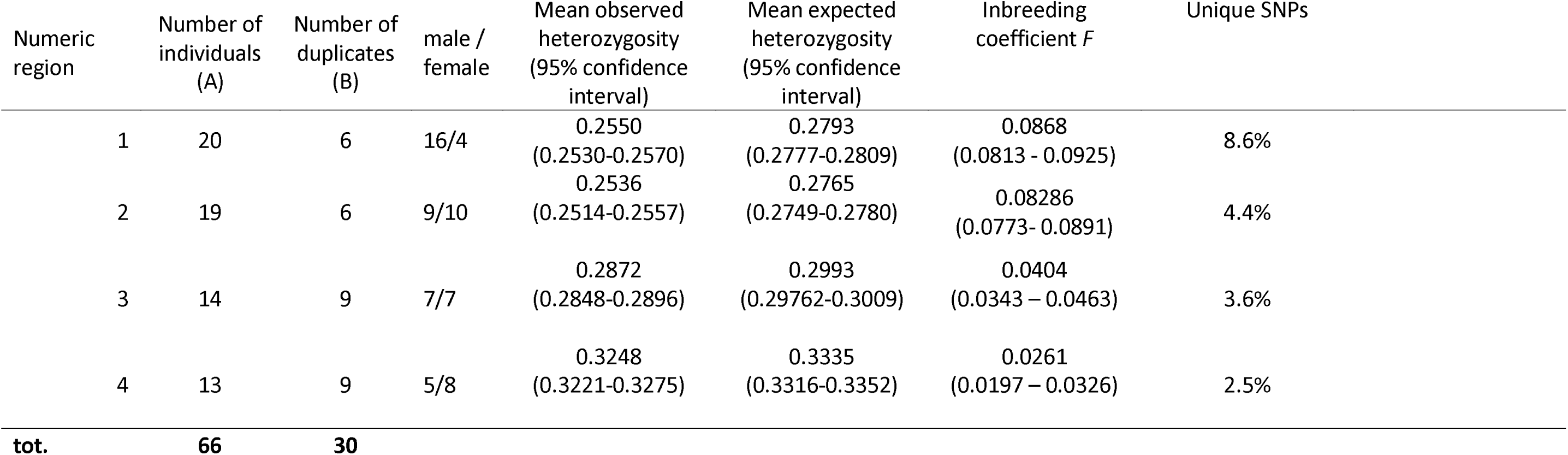
Individuals and samples per region.

| Numeric region | Number of individuals (A) | Number of duplicates (B) | male / female | Mean observed heterozygosity (95% confidence interval) | Mean expected heterozygosity (95% confidence interval) | Inbreeding coefficient *F* | Unique SNPs |
| --- | --- | --- | --- | --- | --- | --- | --- |
| 1 | 20 | 6 | 16/4 | 0.2550<br>(0.2530-0.2570) | 0.2793<br>(0.2777-0.2809) | 0.0868<br>(0.0813 - 0.0925) | 8.6% |
| 2 | 19 | 6 | 9/10 | 0.2536<br>(0.2514-0.2557) | 0.2765<br>(0.2749-0.2780) | 0.08286<br>(0.0773- 0.0891) | 4.4% |
| 3 | 14 | 9 | 7/7 | 0.2872<br>(0.2848-0.2896) | 0.2993<br>(0.29762-0.3009) | 0.0404<br>(0.0343 – 0.0463) | 3.6% |
| 4 | 13 | 9 | 5/8 | 0.3248<br>(0.3221-0.3275) | 0.3335<br>(0.3316-0.3352) | 0.0261<br>(0.0197 – 0.0326) | 2.5% |
| **tot.** | **66** | **30** |  |  |  |  |  |

The amount of DNA in the final solution of each sample was tested and quantified using the Thermo Scientific Nanodrop spectrophotometer, giving a result for the amount of DNA in ng/ μL. Samples were frozen between DNA extraction and analysis.

### Identification and quantification of *P. destructans*

Quantification of *P.destructans* load by qPCR was completed as described previously in Johnson et al. (2015) with the exception of using 1 μl sample in the reaction, Roche Fast Start Essential DNA Probe Master, and a Roche Lightcycler 480 instead of a BioRad iCycler.

### RAD sequencing

We sequenced 66 individuals in total. Duplicate samples from 30 individuals were additionally sequenced to estimate repeatability and error rate of the called genotypes. DNA was prepared for genotyping-by-sequencing using a double digestion RAD-tag method as described in Elshire et al. (2011). PstI-BamHI-digested libraries were prepared by the Center of Evolutionary Applications (University of Turku; see Lemopoulos et al. 2017 and references therein for further details) and sequenced in an Illumina HiSeq2500 run (100 bp single end reads and pooling 96 samples on each lane) at Finnish Functional Genomics Centre (Turku Bioscience).

### Yield comparison samples

As the amount of DNA available may often be very limited in experiments where preserved samples from the rabies laboratory are utilized, we wanted to estimate the effect of initial DNA concentration to the resulting read coverage. We compared the total read coverages of the replicate samples with Pearson’s correlation.

The resulting fastq reads, separated by barcode for each sample, were quality-controlled and low-quality bases trimmed with condetri v. 2.3 with parameter minlen=30 (min. length of a trimmed read) followed by adapter-trim with cutadapt for Illumina universal adapters from the end of the reads (identified with FastQC quality control in some of the samples). Then, reads were mapped against *M. lucifugus* genome (mluc_2.0_assembly_supers.fasta) using bwa mem with parameters -B 3 -O 5 -k 15. After mapping back with BWA mem, we used SAMtools v. 1.4 and the associated bcftools for calling genotypes for SNPs and for filtering SNPs based on minimum of 40% of the samples genotyped, at least eight alternative alleles detected and SNP quality >=20 (bcftools filter). We further filtered the SNPs based on exactly two alleles detected and excluded SNPs with particularly low (≤5) or unusually high (≥125) mean coverage based on visual inspection of the mean coverage distribution.

### Population genetic analysis

For the final SNP dataset, which only included unduplicated samples, we performed principal component analysis implemented in prcomp function of R stats package, and calculated Euclidean genetic coordinates for each individual from PC1 and PC2. The four pre-determined ancestral populations based on the sampling regions of each individual were confirmed by hierarchical clustering of the Euclidean distanced calculated from PC1 and PC2. Individuals clustering to a neighbouring population were assumed to have dispersed from their natal populations and were reassigned for the later analysis. We further investigated the relative contributions of the ancestral populations inhabiting regions 1-4 in the present-day nucleotide using ADMIXTURE (Zhou et al. 2009) analysis for the A-samples, run with four expected populations (parameter K=4) based on prior knowledge of the population structure, and with quasi-Newton convergence acceleration method. We were particularly interested in identifying possible hybrid individuals.

After assigning each individual to the final regions, we calculated Nei’s pairwise *F*_ST_ using HierFSTat v. 0.04-22 pairwise. FST function for A-samples for SNP’s with no missing values. To assess the significance of genetic differentiation, we compared the actual obtained *F*_ST_ estimates to the null distributions of *F*_ST_ values under panmixia, obtained from a hundred random permutations of alleles. Latitude and longitude coordinates of the sampling locations were used to calculate pairwise geographic distances between individuals in kilometres using Haversine method assuming a spherical earth, implemented in function distm in the R package geodist v. 1.5-10.

We estimated Isolation-by-distance with two methods: A Mantel test with complete permutations and a linear model *geographic distance ∼ genetic distance*. We used pairwise *F*_ST_ as a measure for the genetic distances and mean of between-individual distances as geographic coordinates for populations inhabiting each of the four regions. To study the population structure in more detail, we repeated the Isolation-by-distance analysis for the genetic vs. geographic distances from the most extreme individual (sample ID 700) using a Mantel test with all possible permutations, and a linear model to identify individuals with unusually high or low genetic differentiation. We then studied the systematic differentiation of the individuals sampled in different regions by assessing the differences of the residual distributions of each of the four populations from zero using t-tests and Bonferroni-correction of the *P*-values.

We also calculated mean observed heterozygosity within the variable loci in Hardy-Weinberg equilibrium (*FDR*>=0.05) in each of the four study regions and compared the observed values to the mean expected heterozygosities using inbreeding coefficient *F,* calculated as the difference between expected and observed heterozygosity divided by expected heterozygosity (Serre 2006). The 95% confidence intervals for the heterozygosity estimates were found by randomly sampling the variable loci for 1,000 times and extracting the distributions of bootstrap means. The deviations of *F* statistics from zero were detected using single-sample Wilcoxon tests. The significances of the regional differences in the observed and expected heterozygosity distributions, and in the F statistics, were tested both by inspecting the overlaps in the bootstrapped confidence intervals and using analysis of variance and Tukey’s post hoc tests. The significances of within-region differences between the expected and observed heterozygosity were tested both by comparing the bootstrapped confidence intervals and by pairwise t-tests and Bonferroni-corrected p-values. Finally, the fraction of SNPs unique to any one region, and the number of SNPs shared between all regions, were calculated from the observations of variable and non-variable loci within regions.

### Demographic modeling

To formally test whether any of the studied populations experienced secondary contact following glaciation, we implemented a demographic analysis using the software *fastsimcoal2* (Excoffier *et al*. 2013) in combination with the folded site frequency spectra (SFS) calculated from our data using easySFS.py utility (available from https://github.com/isaacovercast/easySFS). Specifically, we evaluated support for two alternative models (Supplementary Figure S6). The first model, representing our null hypothesis of no secondary contact among populations, specified four distinct lineages (region 1, region 2, region 3 and region 4) corresponding to the populations inhabiting the four geographic regions sampled in the present study. These diverged from each other at the time points T1, T2 and T3 as presented in Figure S6a, and exchanged no migrants. Region 1 was used as the lineage from which the other three populations emerged, as this is thought to be the population which is closest to the ancestral *Myotis chiloensis* population. The second model, representing our alternative hypothesis of secondary contact among populations, was identical with the exception that symmetric migration was present between population pairs region 1-region 2 and region 3-region 4, and asymmetric migration was implemented from region 2 to region 3 (Supplementary Figure S6b). In addition to identify the model that was best supported by our data, we also estimated divergence times (T1, T2 and T3) and effective population sizes (region 1, region 2, region 3 and region 4) for the four modeled lineages, as well as migration rates (Mig12, Mig21, Mig34, Mig43 and Mig32).

We performed 50 independent *fastsimcoal2* runs for each model, with 100,000 simulations and 40 cycles of the likelihood maximization algorithm. We then calculated Akaike’s Information Criteria (AIC) from the *fastsimcoal2* runs which yielded the highest maximum likelihood for each model and used these values for model comparison. Finally, we extracted parameter estimates from the best run of the most supported model and calculated 95% confidence intervals based on 100 parametric bootstrap replicates, as described in Excoffier et al. (2013).

### Data availability

The RAD sequencing reads were deposited at NCBI SRA under BioProject ID PRJNA596389.

## RESULTS

### Filtering

Ninety-one of the 96 samples (66 individuals and 30 duplicates) had reads matched with a barcode. Of the obtained genotypes, 54846 were biallelic and used in the later analysis, while we excluded 88882 non-variable (homozygous to alternative) variants and 1708 variants with more than 2 alleles. After filtering, the mean SNP coverage ranged from 0.5 to 257.0 (Supplementary Figure S1). We excluded tags with < 5 or > 125 mean coverage, leaving 47079 tags. The average rate of missing SNPs among the final unique samples was 6.2%, ranging from 0 to 24%.

### Replicate samples

The correlation between input DNA concentration and the resulting mean per-sample read coverage was only moderate (cor = 0.3394, t_61_ = 2.8178, *P* = 0.0065, Supplementary Figure S2 A, Supplementary Figure 3 A-B). In contrast, we found strong and negative association between mean read coverage after sequence assembly and the number of missing genotype calls (cor = -0.9763, t_61_ = -35.223, *P* < 2.2e-16, Figure S2 B). Furthermore, read coverages were very similar between the technical replicate samples (cor = 0.8736, t_26_ = 9.1542, *P* = 1.292e-09, Supplementary figure S2 C), allowing us to omit “B” samples (replicated). The removal of the biological replicates was conducted to minimise the possible SNP calling bias induced by some samples having approximately twice the amount of sequence data compared to the others, if replicate samples had been combined. Identical genotype calls ranged from 85.6% to 96.9% with an average of 94.2% identical genotype calls (Supplementary figure S3 C), depending heavily on read coverage (cor = 0.9275, t_26_ = 12.641, *P* = 1.312e-12 and cor = 0.8484, t_26_ = 8.1716, *P* = 1.186e-08 in “A” and “B” samples, respectively; Supplementary figure 2 E-F). Although the correlation between the initial DNA concentration and identical genotype calls between biological replicates was significant (cor = 0.5990, t_26_ = 3.814, *P* = 0.0007579; Supplementary figure S2 D), this seemed to mainly be due to two outlier observations with both very low concentration and repeatability.

### Identification and quantification of *P. destructans*

Besides our control samples, no samples showed signs of amplification a portion of the multicopy intergenic spacer region of the rRNA gene complex of *P. destructans* by 38 cycles, which is generally considered as a cut-off for the presence of the pathogen DNA in the samples (Muller *et al*. 2013). Therefore, we can conclude that the *M. chiloensis* individuals sampled in this study did not carry *P. destructans*.

### Population genetic analysis

Principal component analysis on 66 individuals and 5538 SNPs without any missing values from non-duplicated samples (label including A, Figure 1 B) revealed a clear structuring of individuals according to the sampling location indicative of strong population structure. Based on hierarchical clustering of the two most important principal components (Supplementary Figure 4), we confirmed the four pre-determined populations based on the natural hierarchical structuring of the data. Based on the clustering, the sub-population assignment of one individual, sample number 679, changed from Biobio to Maule (which was the most common region assignment among the three nearest neighbors for that individual). The estimation of ancestry of each sampled individual by examining the relative contributions of ancestral populations inhabiting regions 1-4 in the present-day revealed particularly pure ancestral lines in the northern and southern parts of the range, with hybridization occurring in the central part of the range (Figure 2.).

**Figure 2.**
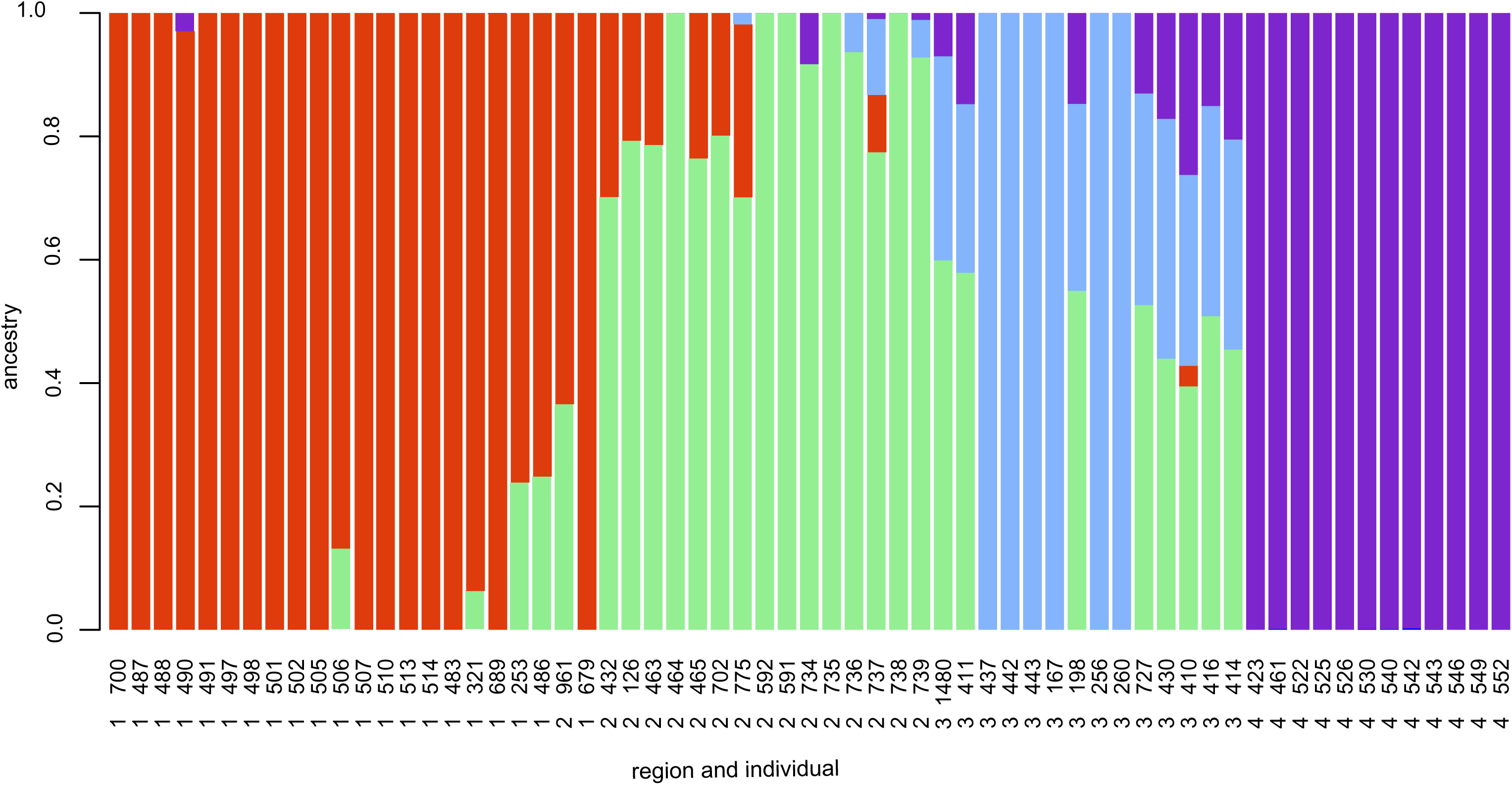
Biogeographical ancestry (admixture) analysis based on nucleotide polymorphisms. Each vertical bar represents an individual, ordered by latitude. Red, green, blue and purple colours indicate the relative genetic contributions of ancestral populations inhabiting regions 1-4, respectively.

Pairwise *F*_ST_-value estimates between the populations sampled from the four geographical locations ranged from 0.04 (between regions 2 and 3) to 0.113 (between the most distant regions 1 and 4, Table 2). All estimated *F*_ST_ values were found significantly larger (*P*<0.01) than the permuted *F*_ST_ distributions, with the 95 % confidence intervals of the null distributions between 0 and 0.0205. Both the Mantel test approach and linear modelling between genetic distances (Euclidean distances calculated from PC1 and PC2) and geographical distances (latitude/longitude coordinates converted to distances in kilometres) strongly suggested that we reject the null hypothesis of geographic and genetic distances being unrelated. For the between-population comparisons with F_ST_, we calculated Mantel statistic r = 0.9497 (*P* = 0.041667) and R-squared estimate of 0.8773 (DF = 1,4, t = 6.063, *P* = 0.00374) (Supplementary figure 5. Also, for the between-individual distances, the observed Mantel statistic r = 0.944 (*P* = 0.001), and R-squared estimate 0.943 (DF = 1,61; t = 31.994; *P* < 2e-16) from the linear modelling suggesting that genetic and geographic distances are strongly positively associated (Figure 3). More detailed exploration of the between-individual linear model revealed, that the distribution of the residuals in regions 1 (DF = 20, t = -3.0679, Bonferroni *P* = 0.024284) and 4 (DF = 11, t = -10.523, Bonferroni *P* = 1.7724e-06) were marginally smaller than 0, while the residuals of individuals in regions 2 (DF = 15; *t* = 3.5896; adj. Bonferroni *P* = 0.002682) and 3 (DF = 13, t = 5.9325, Bonferroni *P* = 0.0001986) were significantly larger than 0 (Figure 3).

**Fig. 3.**
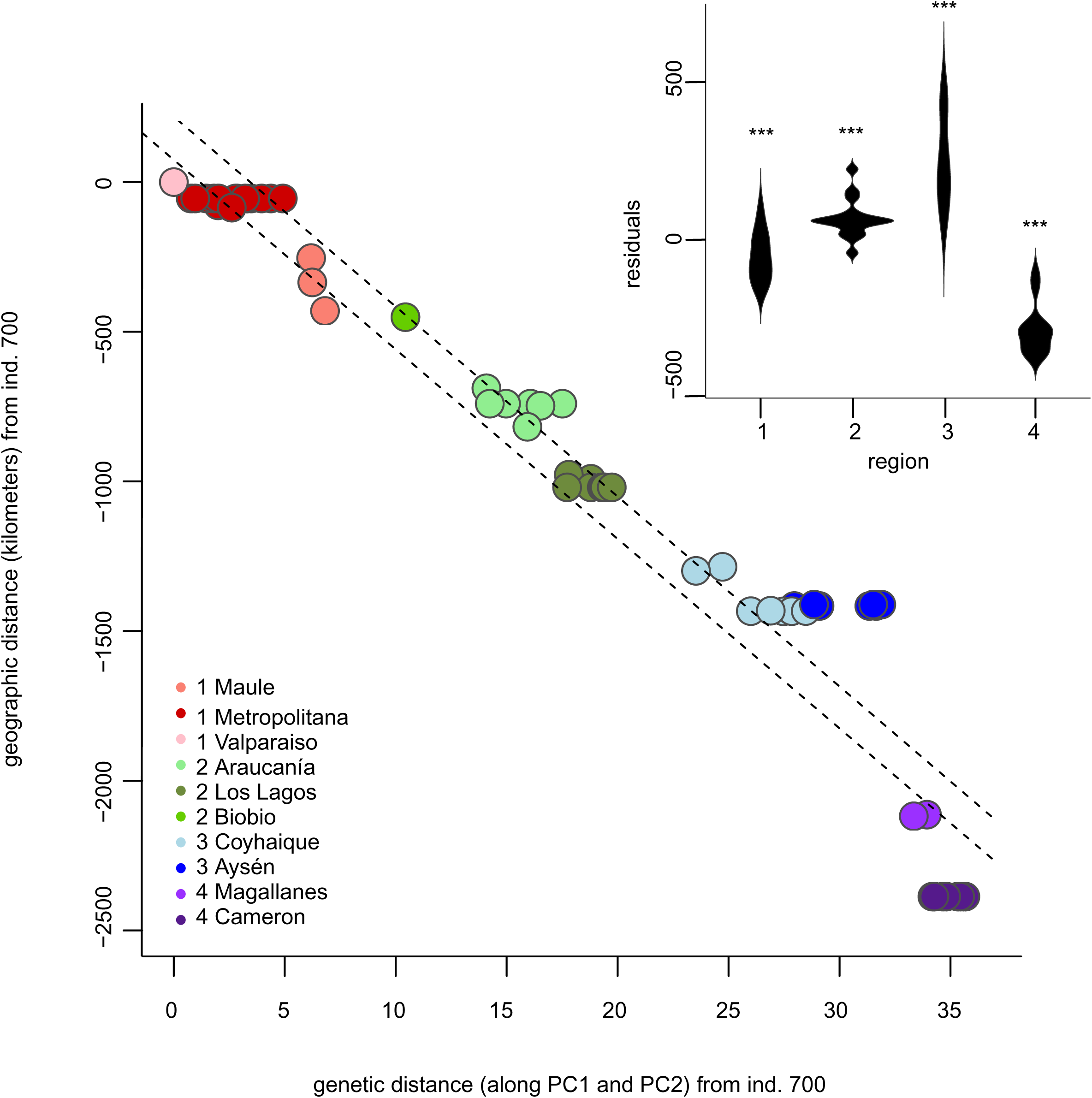
The correlation between geographic and genetic distances between individuals. Geographic distances are measured in kilometres, and genetic distances as Euclidean principal component distances from the reference individual (sample 700). Dashed lines represent 95% confidence interval for the linear model. The violin plot highlights the differences between model residuals for each study region. Residual distributions that differ significantly from zero after Bonferroni correction for multiple testing are marked with * (*P* < 0.05) and *** (*P* < 0.0001).

**Table 2.** Nei’s pairwise F_ST_ and geographic distances (in italics) between populations inhabiting the four geographic regions. The significance level (*P*<0.01) of the F_ST_ statistics is denoted with ***.

| **$F_{ST}$ and mean geographic distance (*km*)** |  |  |  |  |
| --- | --- | --- | --- | --- |
|  | Region 4 | Region 3 | Region 2 | Region 1 |
| Region 4 |  | *970.9* | *1550.3* | *2266.9* |
| Region 3 | 0.075<sup>***</sup> |  | *588.7* | *1326.6* |
| Region 2 | 0.089<sup>***</sup> | 0.041<sup>***</sup> |  | *757.0* |
| Region 1 | 0.113<sup>***</sup> | 0.072<sup>***</sup> | 0.040<sup>***</sup> |  |

Of the total of 47079 SNPs, 43903 were found to be in Hardy-Weinberg equilibrium. Analyses of variance revealed statistically significant differences between populations in observed heterozygosities [*F*(3,106450)=760.6, *P*<2e-16], in expected heterozygosities [*F*(3,106450)=871.4, *P*<2e-16], and in the F statistics [F(3,106450)=55.29, P<2e-16]. Both based on confidence intervals and pairwise comparisons, observed and expected heterozygosity distributions were consistently higher (non-overlapping 95% confidence intervals and adjusted *P*<0.05) in the southern regions than in north, except for the two northernmost regions (region 1 vs. region 2), where a statistically significant difference was not observed (Table 1, Table S2). Similarly, observed heterozygosities were consistently lower than those estimated from allele frequencies (non-overlapping 95% confidence intervals and adjusted *P*<0.05), which may be caused by within-region population structure (Table 1, Table S2). Finally, pairwise comparisons of the inbreeding coefficient distributions indicated that the excess of homozygotes was greater in the north than in the south when compared to neutral expectation based on allele frequencies. This was supported with statistically significant differences (non-overlapping confidence intervals and adjusted *P*<0.05) observed in all comparisons except in region pairs 2 and 3; and 4 and 5 (Table 1, Table S2). This indicated that the northern populations are inbreeding more than the southern populations (Table 1, Table S2).

A large proportion of SNPs, 25.3%, were polymorphic in all four regions. The fraction of SNPs unique only to one region decreased from going north to south: while 8.6% of the SNPs were unique to region 1, only 2.5% unique SNPs were found in region 4.

We did not find support for secondary contact using demographic modelling with Fastsimcoal2. Instead, the null model that included no migration among populations (Figure S6a) received the highest AIC support (Table S3). Parameter priors and estimates, together with their 95% confidence intervals, from the best model are reported in Table S4.

## DISCUSSION

Our results present the first assessment of population structure in the widely distributed bat species, *M. chiloensis*, using individual-based approach with genome-wide markers. We found that geographic distance within the range of the species are reflected in its population structure. Although we found a clear and robust population structure among sampling sites, population structure is correlated with geographical distance, even though populations are separated by ice fields, mountain ranges and stretches of open water. This has implications for the protection of populations that may be susceptible to WNS. Our results also show highest genetic variability in the species is at the southern extent of its range.

Strong population structure is rarely seen in bats, even across large geographical scales in genera such as *Myotis*, with shorter dispersal distances (Castella *et al*. 2000; Atterby *et al*. 2010; Laine *et al*. 2013). In fact, geographical distance often correlates significantly with genetic distance in bats. This is partially due to autumn migration and swarming behaviour in *Myotis* species, which brings together bats from broad geographic areas to breed and promote recombination (Burns *et al*. 2014; Burns and Broders 2015). Furthermore, powered flight allows effective dispersal, which is often male biased (Arnold 2007; Angell *et al*. 2013). This is reflected in low fixation indices in widespread bat species, such as *M. daubentonii*, where individuals separated by thousands of kilometres in Europe show low fixation indexes (Laine *et al*. 2013). While fixation indices cannot be compared directly across species, especially when different methodological approaches are used, they do give an indication of the connectivity of populations.

Although geographic and genetic distances were found to have a strong positive correlation in our study, we can concur that the southernmost population shows a higher degree of isolation compared to the geographic distance to its closest comparative population to the north. An F_ST_ of 0.075 between regions 3 and 4 using whole-genome data is already higher than F_ST_ values recorded for *M. daubentonii* across Europe using microsatellites (Laine *et al*. 2013), and in our data, the geographic distance is only c. 1000 km. Our data, with tens of thousands of SNP’s also allowed a more precise estimate of fixation compared to a handful of microsatellites. However, due to a limited number of individuals, it did not allow us to examine dispersal as a function of sex, which in bats is often a male driven function (Arnold 2007; Laine *et al*. 2013; Angell *et al*. 2013). Our sampling may also have missed some connecting populations in between regions 3 and 4, which could for instance be located on the eastern slopes of the Andes. However, our test for relative contributions of ancestral populations reveals bats in region 4, in the Magallanes and Tierra del Fuego, have no mixing of ancestral populations with the other regions.

In the northern hemisphere, approximately 18000 years ago, at the end of the Late Pleistocene, the ice sheets began to recede as the global climate became warmer. The biota migrated northwards following their optimal environments (Huntley and Webb 1989). This expansion of refugial populations has been associated with genetic variation decreasing south-to-north in some species: a trend attributed to a series of bottlenecks when the biota spread from the leading edge of the refugial population, leading to a loss of alleles and decreasing genetic diversity (Hewitt 1996, 1999). Contrary to what one could expect based on these latitudinal shifts in diversity in the northern latitude, genetic diversity in *M. chiloensis* appears to increase with increasing latitude, from north to south. We presumed this counterintuitive pattern of heterozygosity within the species may be related to the glaciation history of South America, where populations isolated by glacial events could have been able to hybridize. For instance, in some terrestrial European and Scandinavian vertebrates, the intraspecific genealogical lineages, which formed in separate refugia, were found to have come to secondary contact in the Fennoscandian area (Tegelström 1987; Jaarola and Tegelström 1995; Nesbø *et al*. 1999; Knopp and Merilä 2009).

*Myotis chiloensis* is described as a vicariant species with respect to other closely related *Myotis* species (*M. albescens, M. nigricans, M. levis*) from South America, Ruedi et al (2013) estimated this isolation event at 5.5 My in late Miocene. This same time period saw the beginning of a number of glaciations events in Patagonia with variable intensity and duration (Rabassa *et al*. 2011). Glacial episodes isolated the Patagonian forest from around middle Miocene well into the late Quaternary, the Last Glacial Maximum (Rabassa *et al*. 2011, Supplemental Figure S7). During the glaciations, the forests on the Pacific coast were most like completely suppressed, possibly with isolated small refugia. On the Atlantic side, the forest was fragmented from 36°S southwards (Sérsic *et al*. 2011). Finally, in Tierra del Fuego the forest was probably displaced towards the current submarine shelf (Ponce *et al*. 2011). As the ice retreated refugial populations may have come into secondary contact in the southern part of current range, which could explain the high heterozygosity as well as the small fraction of unique SNP’s of these populations. However, our coalescence analysis rejected the secondary contact model, favouring the null model suggesting our study populations are still largely separated after the Last Glacial Maximum. Indeed, the F_ST_-values are high, suggesting isolation of the populations. By contrast, our analysis for the relative contributions of ancestral populations suggests mixing of populations. One potential explanation for this apparent discrepancy is that we derived Site Frequency Spectrum from a rather small number of individuals, which may carry a signal of migration that is not strong enough to allow a model that includes secondary contact to be favoured over a simpler model without migration.

The spread of *P. destructans* via one host to another was very rapid in North America (Blehert *et al*. 2009; Frick *et al*. 2010). This was facilitated in part by the ecology of North American cave-hibernating bats and the availability of suitable environment for the fungus to propagate: limestone caves found throughout the Appalachian region in eastern North America (Lorch *et al*. 2013). Furthermore, as a consequence of down-regulation of metabolism during extended torpor bouts, attempted immune responses fall short, and may even contribute to mortality in hosts infected with *P. destructans* (Field *et al*. 2015; Lilley *et al*. 2017, 2019). In addition to these, the massive population declines associated with WNS in affected species (Turner *et al*. 2011) were magnified by the panmictic population structure across eastern North America in the most affected species, *M. lucifugus* (Miller-Butterworth *et al*. 2014; Vonhof *et al*. 2015).

Our results for *M. chiloensis* from austral South America suggest the southernmost population in region 4, may be less likely to be infected via their continental conspecifics, because of reduced contact between the populations. Our results indicated no mixing of ancestry in the southernmost individuals in our study, suggesting the *M. chiloensis* on Tierra del Fuego are isolated from their mainland counterparts. To our knowledge, this is also the only population to use extended torpor, a prerequisite for the propagation of *P. destructans* and the onset of WNS (Ossa *et al*. Submitted for publication). Tierra del Fuego, the southern tip of Patagonia and the continent of South America, experiences extended low winter temperatures comparable to areas in North America where WNS is manifested. Our genetic analysis for the presence of *P. destructans* on the sampled bats suggests the fungal pathogen does not exist within the distribution range of our focal species at present. Even if the fungus were to enter the region, variability in host behavior and environmental characteristics may be the primary factors protecting hosts from the pathology related to WNS (Zukal *et al*. 2014, 2016). Most strikingly large cave hibernacula with suitable, stable environmental conditions favouring the environmental persistence of the pathogen in the absence of the hosts, are very scarce and separated by hundreds of kilometers in most of the southern range of *M. chiloensis*.

The observed distribution of *M. chiloensis* is vast, covering a range of forested habitats from arid Sclerophyllous to sub-Antartic (Ossa and Rodriguez-San Pedro 2015). In this respect, taking into consideration our results on clear population segregation begs to propose the question on the species status of *M. chiloensis* as a whole. Indeed, *M. chiloensis* also appears to vary phenotypically along its distribution range (Mann 1978; Ossa 2016). It has been proposed that the species was composed of three sub-species: *M. ch. atacamensis* which is presently known as *M. atacamensis; M. ch. arescens* from 29°S to 39°S; and *M. ch. chiloensis* from 39°S to 53°S (Mann 1978). That classification was due according to their coat colour changes in relation to exposure to solar radiation and ambient temperature, which is correlated to latitude (Budyko 1969) and levels of precipitation in their habitat. They vary from a lighter pelage to a dark brown colour on a gradient from the northern part of their range to the south (Galaz *et al*. 2006). This promotes the theory, isolation by adaptation, as a driver of population genetic structure. The genetic adaptations of an individual to their local environment separates populations and leads to a reduced gene flow (Orsini *et al*. 2013). However, the isolation by adaptation theory negates the fact that there is a possibility of a barrier so that gene flow is inhibited by climate or adaptations to the local environment. It is possible to therefore state that isolation-by-dispersal limitation, and moreover isolation-by-distance, are the more probable causes of the observed results in the present study. Indeed, the reluctance of the species to cross barriers, for instance the Andes, can clearly be seen by examining population 3, where six individuals sampled from Puerto Aysén (437, 442, 443, 167, 256, 260) are visibly isolated on the PCA plot form individuals on the other side of the Andes, under 150 km away. This also depicts the fine scale resolution our individual-based SNP-based approach allows. However, further studies should focus deeper on the taxon status of different population of the species currently recognized as *M. chiloensis*.

The results highlight the importance to assess the population structure which may limit the spread of white-nose syndrome disease. Whether *P. destructans* or another epizootic in the future could spread depends largely on the population structure and connectedness of hosts (Lilley *et al*. 2018).

## ACKNOWLEDGEMENTS

We thank the Rufford Foundation (Rufford Small Grant 10502-1 and 23042-2), H2020 Marie Skłodowska-Curie Actions (706196), and Ohio University Research Council for funding the work. We thank Servicio Agricola y ganadero Diproren for the capture permits in Tierra del Fuego (Res Ex: 1253/2016 and 4924/2017), Juan Carlos Aravena from the Instituto de la Patagonia for his help, the personnel from WCS Chile for allowing us to conduct research at Karukinka Natural Reserve as well as for their help with field work. We thank Michelle Lineros and Tania Gatica from the National Health Institute for their help to obtain the tissue samples. We thank Austin Waag for assistance with field work.

