## Supplementary figures and tables for "Population connectivity predicts vulnerability to white-nose syndrome in the Chilean myotis (*Myotis chiloensis*) - a genomics approach"

**Table S1.**

| **Sample ID** | **region ID** | **Date** | **Live sampled** | **Latitude** | **Longitude** | **Locality** | **Geographical Region** | **Age** | **Sex** | **Weight** | **Forearm Length** |
| --- | --- | --- | --- | --- | --- | --- | --- | --- | --- | --- | --- |
| 253 | 1 | 19/01/2017 |  | -34.966525 | -71.25523393 | Curico | Maule | A | M |  | 37.4 |
| 321 | 1 | 23/01/2017 |  | -33.398303 | -70.51509469 | Las Condes | Metropolitana | A | F |  | 38.7 |
| 483 | 1 | 15/02/2017 |  | -33.220534 | -70.71641614 | Colina | Metropolitana | A | M |  | 37 |
| 486 | 1 | 16/02/2017 |  | -35.684996 | -71.41426684 | Colbún | Maule | A | F |  | 38 |
| 487 | 1 | 07/11/2017 | x | -33.186147 | -70.94819205 | Lampa | Metropolitana | A | M | 5.6 | 36.8 |
| 488 | 1 | 07/11/2017 | x | -33.186147 | -70.94819205 | Lampa | Metropolitana | A | M | 6 | 35.7 |
| 490 | 1 | 07/11/2017 | x | -33.186147 | -70.94819205 | Lampa | Metropolitana | A | M | 5.4 | 35.3 |
| 491 | 1 | 07/11/2017 | x | -33.186147 | -70.94819205 | Lampa | Metropolitana | A | M | 5.8 | 37 |
| 497 | 1 | 08/11/2017 | x | -33.186147 | -70.94819205 | Lampa | Metropolitana | A | M | 6.4 | 38.2 |
| 498 | 1 | 08/11/2017 | x | -33.186147 | -70.94819205 | Lampa | Metropolitana | A | M | 6.4 | 37.8 |
| 501 | 1 | 08/11/2017 | x | -33.186147 | -70.94819205 | Lampa | Metropolitana | A | M | 6 | 39.5 |
| 502 | 1 | 08/11/2017 | x | -33.186147 | -70.94819205 | Lampa | Metropolitana | A | M | 7 | 36.4 |
| 505 | 1 | 08/11/2017 | x | -33.186147 | -70.94819205 | Lampa | Metropolitana | A | M |  |  |
| 506 | 1 | 08/11/2017 | x | -33.186147 | -70.94819205 | Lampa | Metropolitana | A | M |  |  |
| 507 | 1 | 08/11/2017 | x | -33.186147 | -70.94819205 | Lampa | Metropolitana | A | M |  |  |
| 510 | 1 | 13/11/2017 | x | -33.186147 | -70.94819205 | Lampa | Metropolitana | A | M | 6.6 | 35.4 |
| 513 | 1 | 13/11/2017 | x | -33.186147 | -70.94819205 | Lampa | Metropolitana | A | M | 6 | 36.5 |
| 514 | 1 | 13/11/2017 | x | -33.186147 | -70.94819205 | Lampa | Metropolitana | A | M | 6.6 | 36.4 |
| 689 | 1 | 20/03/2017 |  | -33.490726 | -70.6544068 | San Miguel | Metropolitana | A | F |  | 37 |
| 700 | 1 | 23/03/2017 |  | -32.726856 | -70.73995691 | San Felipe | Valparaiso | A | M |  | 36 |
| 126 | 2 | 11/01/2017 |  | -39.285082 | -72.09628668 | Villarrica | Araucanía | A | F |  | 36.2 |
| 432 | 2 | 29/12/2016 |  | -38.727988 | -72.59287963 | Temuco | Araucanía | J | F | 4.4 | 37.1 |
| 463 | 2 | 10/02/2017 |  | -39.285082 | -72.09628668 | Villarrica | Araucanía | A | F |  | 38.6 |
| 464 | 2 | 10/02/2017 |  | -39.285082 | -72.09628668 | Villarrica | Araucanía | A | F |  | 39.8 |
| 465 | 2 | 10/02/2017 |  | -39.285082 | -72.09628668 | Villarrica | Araucanía | A | F |  | 38.7 |
| 590 | 2 | 02/03/2017 |  | -41.450413 | -72.97563916 | Puerto Montt | Los Lagos | A | M |  | 36.4 |
| 591 | 2 | 02/03/2017 |  | -41.450413 | -72.97563916 | Puerto Montt | Los Lagos | A | M |  | 37.7 |
| 592 | 2 | 02/03/2017 |  | -41.330658 | -72.94001505 | Puerto Varas | Los Lagos | A | M |  | 36.6 |
| 679 | 2 |  |  | -36.535425 | -71.549531 | (San Fabian) Curico * | (Biobio) Maule * |  | F |  |  |
| 702 | 2 | 23/03/2017 |  | -39.332327 | -72.21733907 | Villarrica | Araucanía | A | M |  | 36.8 |
| 734 | 2 | 31/03/2017 |  | -41.600315 | -73.59914542 | Maullin | Los Lagos | A | M |  | 37.6 |
| 735 | 2 | 31/03/2017 |  | -41.600315 | -73.59914542 | Maullin | Los Lagos | A | F |  | 36.4 |
| 736 | 2 | 31/03/2017 |  | -41.600315 | -73.59914542 | Maullin | Los Lagos | A | M |  | 36.4 |
| 737 | 2 | 31/03/2017 |  | -41.600315 | -73.59914542 | Maullin | Los Lagos | A | M |  | 38 |
| 738 | 2 | 31/03/2017 |  | -41.600315 | -73.59914542 | Maullin | Los Lagos | A | F |  | 38 |
| 739 | 2 | 31/03/2017 |  | -41.600315 | -73.59914542 | Maullin | Los Lagos | A | F |  | 38.5 |
| 741 | 2 | 31/03/2017 |  | -41.600315 | -73.59914542 | Maullin | Los Lagos | A | F |  | 37.1 |
| 775 | 2 | 06/04/2017 |  | -39.789315 | -73.23833125 | Valdivia | Los Rios | A | M |  | 37.9 |
| 961 | 2 | 05/06/2017 |  | -36.533333 | -72.43398946 | San Ignacio | Biobio | A | M |  | 36.6 |
| 167 | 3 | 13/01/2017 |  | -45.354771 | -72.70681106 | Puerto Aysen | Aysén | A | M |  | 34.2 |
| 198 | 3 | 17/01/2017 |  | -45.354771 | -72.70681106 | Puerto Aysen | Aysén | A | F |  | 36.2 |
| 256 | 3 | 19/01/2017 |  | -45.354771 | -72.70681106 | Puerto Aysen | Aysén | A | M |  | 36.8 |
| 260 | 3 | 19/01/2017 |  | -45.354771 | -72.70681106 | Puerto Aysen | Aysén | A | M |  | 33.9 |
| 410 | 3 | 18/12/2014 |  | -45.561848 | -72.04543466 | Coyhaique | Aysén | A | F | 7.9 | 38.9 |
| 411 | 3 | 03/02/2015 |  | -44.305737 | -72.56037601 | Coyhaique | Aysén | A | F | 7.6 | 38.5 |
| 414 | 3 | 05/03/2015 |  | -45.566087 | -72.06953519 | Coyhaique | Aysén | A | F | 7.1 | 40.3 |
| 416 | 3 | 23/12/2015 |  | -45.561848 | -72.04543466 | Coyhaique | Aysén | A | F | 8.2 | 38.8 |
| 430 | 3 | 03/02/2017 |  | -45.559139 | -72.03808095 | Coyhaique | Aysén | A | F |  | 38.9 |
| 437 | 3 | 13/01/2017 |  | -45.312844 | -72.70248186 | Puerto Aysen | Aysén | J | M | 5.8 | 34.6 |
| 442 | 3 | 19/01/2017 |  | -45.312844 | -72.70248186 | Puerto Aysen | Aysén | J | M | 6.2 | 36.8 |
| 443 | 3 | 19/01/2017 |  | -45.312844 | -72.70248186 | Puerto Aysen | Aysén | J | M | 6.2 | 34.1 |
| 727 | 3 | 29/03/2017 |  | -45.54847 | -72.02953319 | Coyhaique | Aysén | A | M |  | 40.1 |
| 1480 | 3 | 11/10/2017 |  | -44.238521 | -71.851127 | Lago Verde | Aysén | A | F |  | 39.9 |
| 412 | 4 | 18/02/2015 |  | -51.71095 | -72.51273744 | Puerto Natales | Magallanes | A | M | 6.2 | 39.5 |
| 423 | 4 | 10/03/2016 |  | -51.71095 | -72.51273744 | Puerto Natales | Magallanes | A | F |  | 38.8 |
| 461 | 4 | 10/02/2017 |  | -51.71095 | -72.51273744 | Puerto Natales | Magallanes | A | F |  | 40 |
| 522 | 4 | 05/12/2017 | x | -54.116965 | -68.70559422 | Cameron | Magallanes | A | F | 8.8 | 37.8 |
| 525 | 4 | 05/12/2017 | x | -54.116965 | -68.70559422 | Cameron | Magallanes | A | F | 11 | 39.6 |
| 526 | 4 | 05/12/2017 | x | -54.116965 | -68.70559422 | Cameron | Magallanes | A | M | 8 | 38.8 |
| 530 | 4 | 06/12/2017 | x | -54.116965 | -68.70559422 | Cameron | Magallanes | A | F | 11 | 39.1 |
| 540 | 4 | 07/12/2017 | x | -54.116965 | -68.70559422 | Cameron | Magallanes | A | F | 8.8 | 39.6 |
| 542 | 4 | 08/12/2017 | x | -54.116965 | -68.70559422 | Cameron | Magallanes | A | M | 7 | 32.6 |
| 543 | 4 | 08/12/2017 | x | -54.116965 | -68.70559422 | Cameron | Magallanes | A | M | 7 | 37.5 |
| 546 | 4 | 08/12/2017 | x | -54.116965 | -68.70559422 | Cameron | Magallanes | A | F | 11.9 | 38.8 |
| 549 | 4 | 08/12/2017 | x | -54.116965 | -68.70559422 | Cameron | Magallanes | A | M | 7.8 | 38 |
| 552 | 4 | 10/12/2017 | x | -54.116965 | -68.70559422 | Cameron | Magallanes | A | F | 10 | 36.9 |

Samples used for study (* reassigned based on hierarchical clustering)

**Table S2.**

|  | **Regions** | **difference in means** | **lower end point of the interval** | **upper end point of the interval** | **Adjusted *P*** |
| --- | --- | --- | --- | --- | --- |
| observed heterozygosity | 2-1 | -0.00137824 | -0.005431822 | 0.002675 | 0.818603 |
|  | 3-2 | 0.032175391 | 0.028060638 | 0.03629 | 0 |
|  | 4-1 | 0.069734096 | 0.065338757 | 0.074129 | 0 |
|  | 3-2 | 0.033553631 | 0.02947549 | 0.037632 | 0 |
|  | 4-2 | 0.071112335 | 0.066751253 | 0.075473 | 0 |
|  | 4-3 | 0.037558705 | 0.033140707 | 0.041977 | 0 |
| expected heterozygosity | 2-1 | -0.00275967 | -0.005704086 | 0.000185 | 0.075577 |
|  | 3-2 | 0.020019459 | 0.017030612 | 0.023008 | 0 |
|  | 4-1 | 0.054174112 | 0.050981457 | 0.057367 | 0 |
|  | 3-2 | 0.02277913 | 0.019816879 | 0.025741 | 0 |
|  | 4-2 | 0.056933784 | 0.053766012 | 0.060102 | 0 |
|  | 4-3 | 0.034154654 | 0.03094554 | 0.037364 | 0 |
| F statistic | 2-1 | -0.005 | -0.01329632 | 0.003563 | 0.447614 |
|  | 3-2 | -0.030 | -0.03871632 | -0.021603 | 0 |
|  | 4-1 | -0.037 | -0.04592906 | -0.027649 | 0 |
|  | 3-2 | -0.025 | -0.03377329 | -0.016812 | 0 |
|  | 4-2 | -0.032 | -0.04099092 | -0.022853 | 0 |
|  | 4-3 | -0.007 | -0.01581647 | 0.002558 | 0.248283 |
| observed vs. expected heterozygosity within a region | 1 | 0.0242 | 0.02269657 | 0.025791 | 8.80E-16 |
|  | 2 | 0.0229 | 0.02126347 | 0.024461 | 8.80E-16 |
|  | 3 | 0.0121 | 0.01028109 | 0.013895 | 8.80E-16 |
|  | 4 | 0.0087 | 0.006466995 | 0.010901 | 6.74E-14 |

Pairwise differences between the genome-wide distributions of observed heterozygosity, expected heterozygosity and *F* statistics, estimated for the regions inhabited by *M. chiloensis* populations; and differences between the distributions of observed and expected heterozygosity within regions. Tukey’s post-hoc test was used to detect the significant differences between the pairwise comparisons of the observed and expected heterozygosity and *F* statistics. Pairwise t-tests were used to compare the distributions of observed and expected heterozygosities within each of the regions. Tukey’s adjusted *P*-values and Bonferroni-corrected *P*-values are reported to correct for multiple comparisons.

**Table S3.**

| **Model** | **Maximum ln(likelihood)** | **Parameters estimated** | **AIC** |
| --- | --- | --- | --- |
| null | -1506.5 | 7 | 6951.6 |
| secondary contact | -1616.7 | 12 | 7469.0 |

Relative maximum ln-transformed likelihoods, given as the best values among 50 independent runs for each demographic model using Fastsimcoal2; number of estimated parameters for and AIC values for the two demographic models tested.

**Table S4.**

| **Parameter** | **Priors** | | **Estimates for the best model** | | |
| --- | --- | --- | --- | --- | --- |
| **Effective population size (haploid)** | **"secondary contact" -model** | **"null" model** | **estimate** | **95% CI lower** | **95% CI upper** |
| Pop1 | 5000-35000 | 5000-35000 | 5061 | 5051 | 5083 |
| Pop2 | 5000-35000 | 5000-35000 | 5052 | 5068 | 5100 |
| Pop3 | 5000-35000 | 5000-35000 | 5069 | 5051 | 5099 |
| Pop4 | 5000-35000 | 5000-35000 | 5051 | 5069 | 5099 |
| **Divergence time** |  |  |  |  |  |
| T1 | 4425-4750 | 4425-4750 | 4482 | 4427 | 4512 |
| T2 | 7000-12000 | 7000-12000 | 7091 | 7019 | 7128 |
| T3 | 15000-30000 | 15000-30000 | 15188 | 15045 | 15270 |
| **Migration** |  |  |  |  |  |
| MIG12 | 1e-5 - 1e-2 | - | - | - | - |
| MIG43 | 1e-5 - 1e-2 | - | - | - | - |
| MIG21 | 1e-5 - 1e-2 | - | - | - | - |
| MIG34 | 1e-5 - 1e-2 | - | - | - | - |

The priors for all estimated parameters for each demographic model, and parameter estimates from the best null model, with associated 95% confidence intervals. See supplementary figure 6 for parameter definitions.

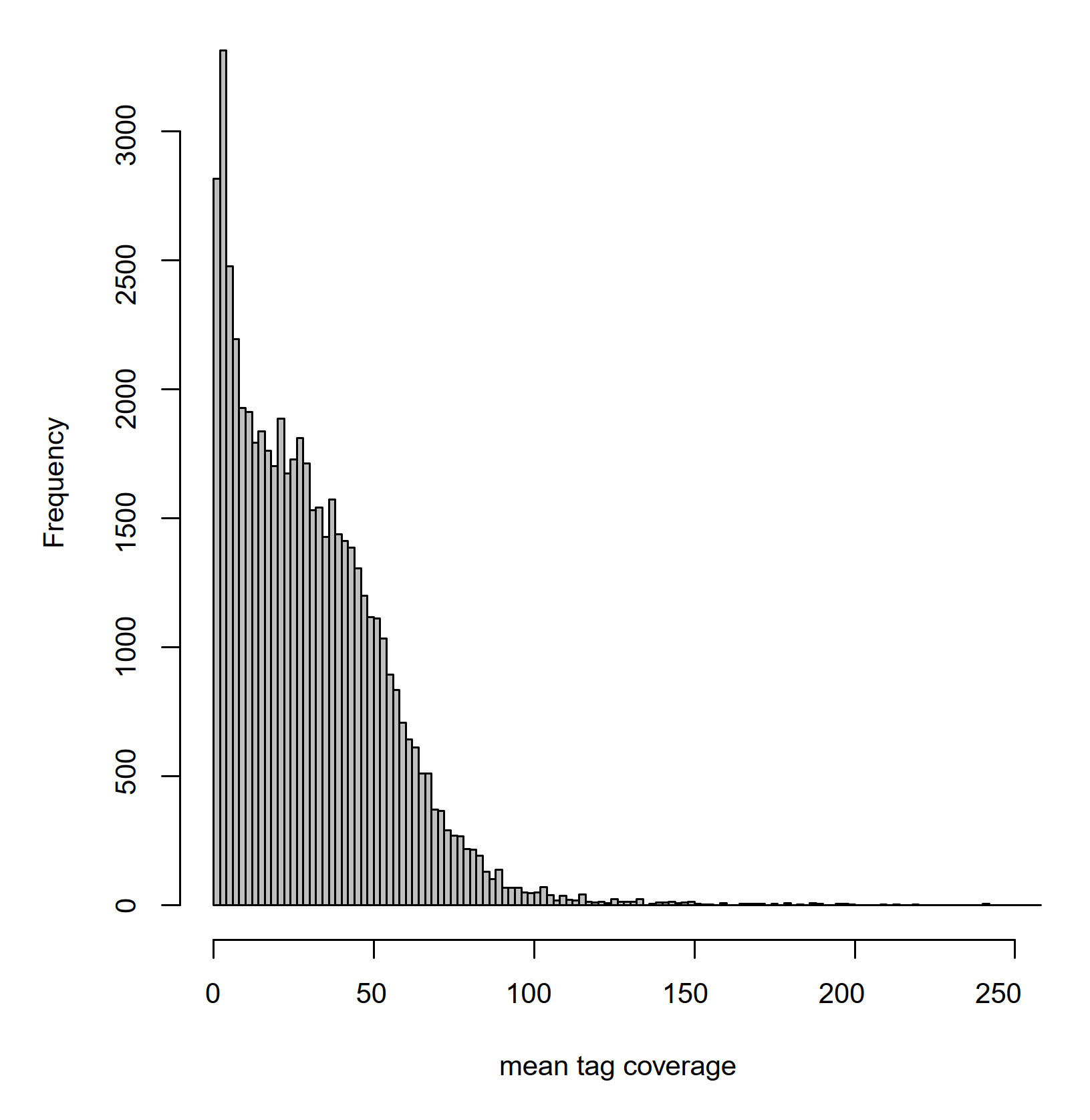

**Figure S1.** The distribution of mean per-individual SNP coverages.

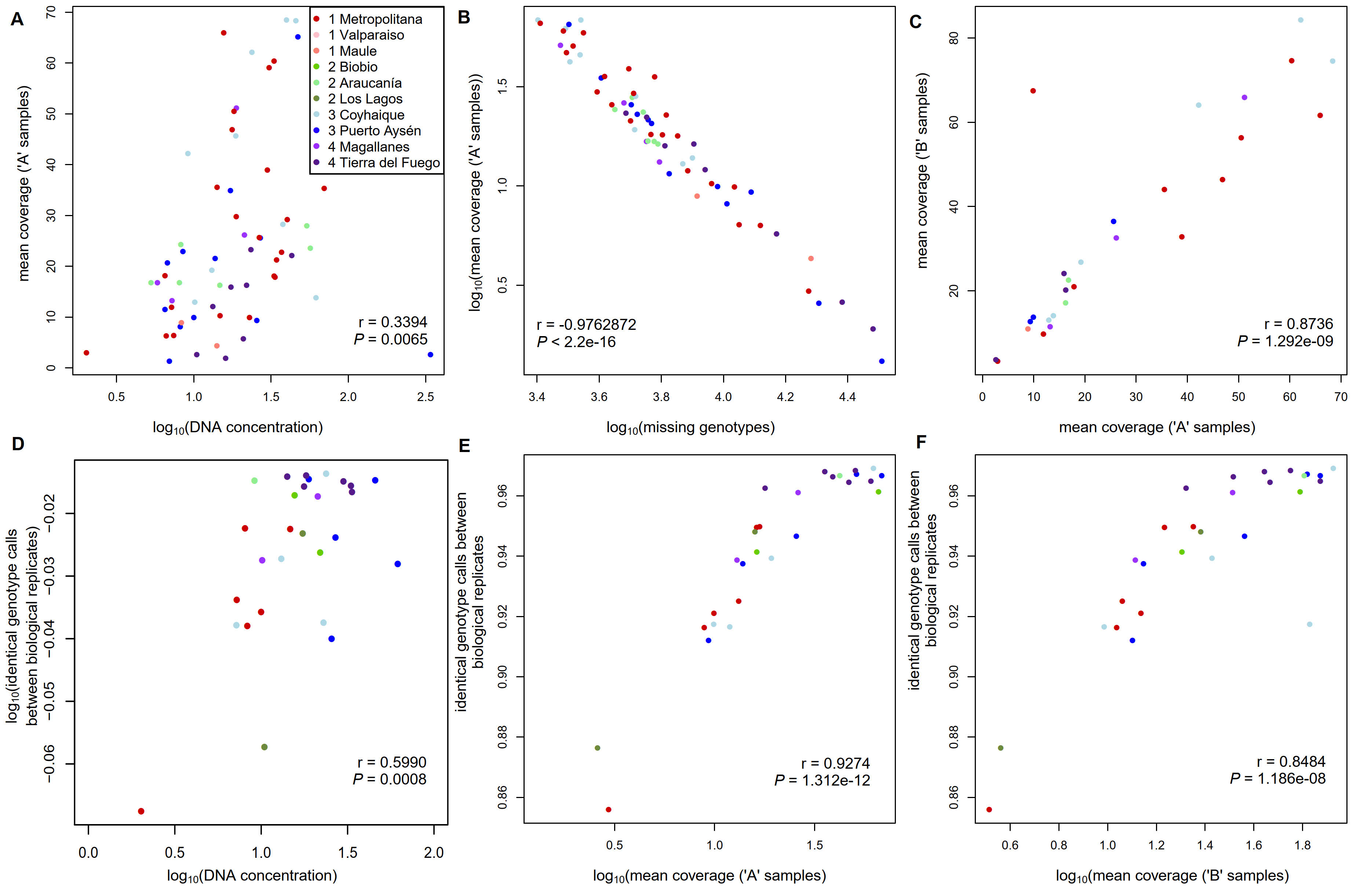

**Figure S2.** A-F. Correlation between DNA concentrations and the resulting per-sample coverages (A); correlation between per-sample coverages and the resulting numbers of missing genotypes (B); correlation between the coverages of biological replicates (“A”- and “B”-samples, C); correlation between identical genotype calls between biological replicates and DNA concentrations (D), and per-sample coverages of “A” samples (E), and per-sample coverages of “B” samples (F).

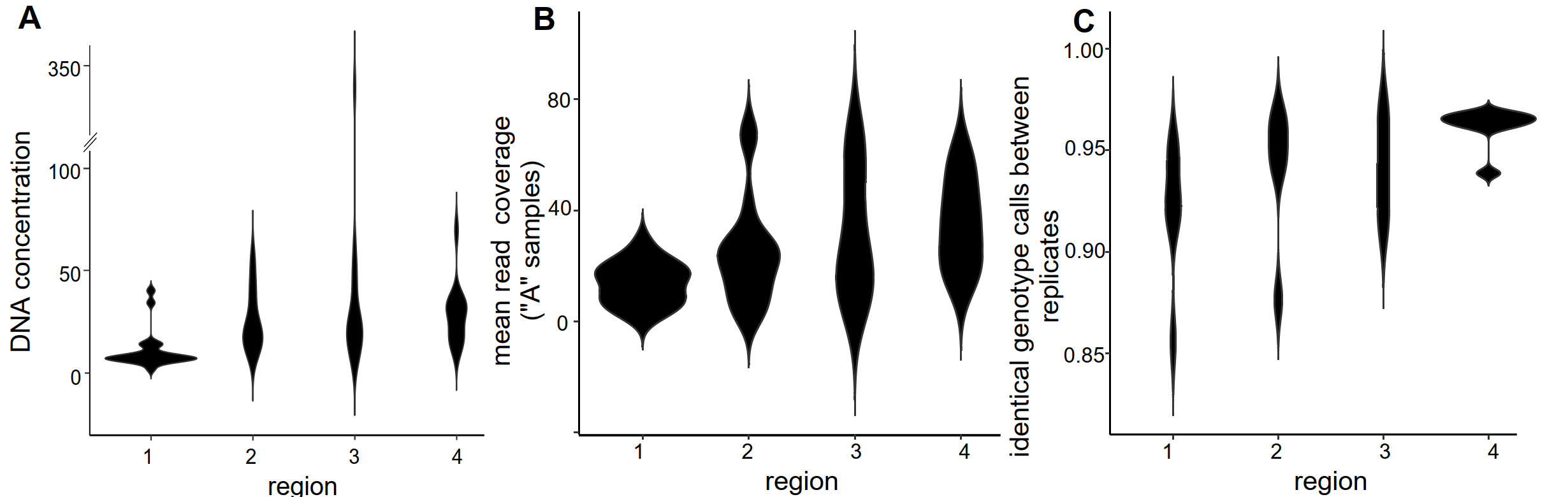

**Figure S3.** The differences in the variation of DNA concentrations (A), in the resulting read coverages after sequence assembly (in “A” samples, B), and finally in the identical genotype calls between biological replicates (C) in regions 1-4.

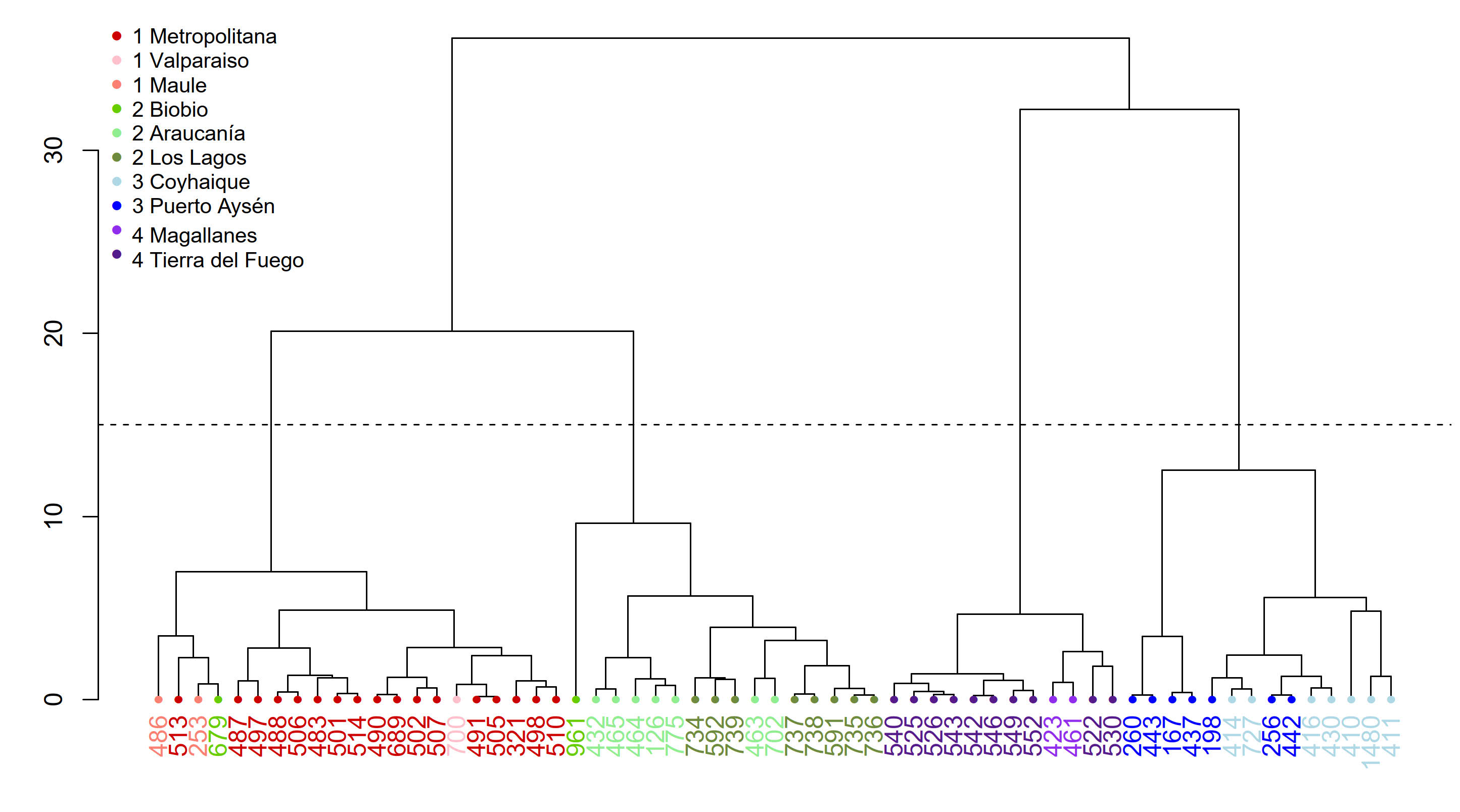

**Figure S4.** Hierarchical clustering based on the pairwise Euclidean distances, calculated from the two most influential principal components between the sampled individuals (‘A’ samples) can be used to naturally divide the data into four clusters (dashed line), except for individual 679, which has probably dispersed from the parental population.

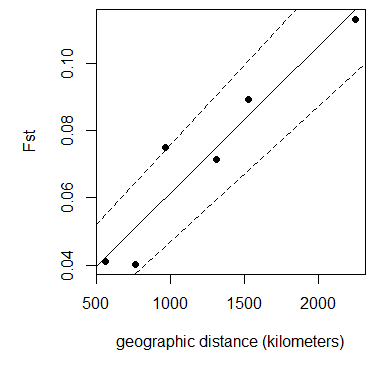

**Figure S5.** Linear model (solid line) and 95% confidence intervals (dashed lines) between pairwise F_ST_ estimates and the mean geographic distance between population pairs.

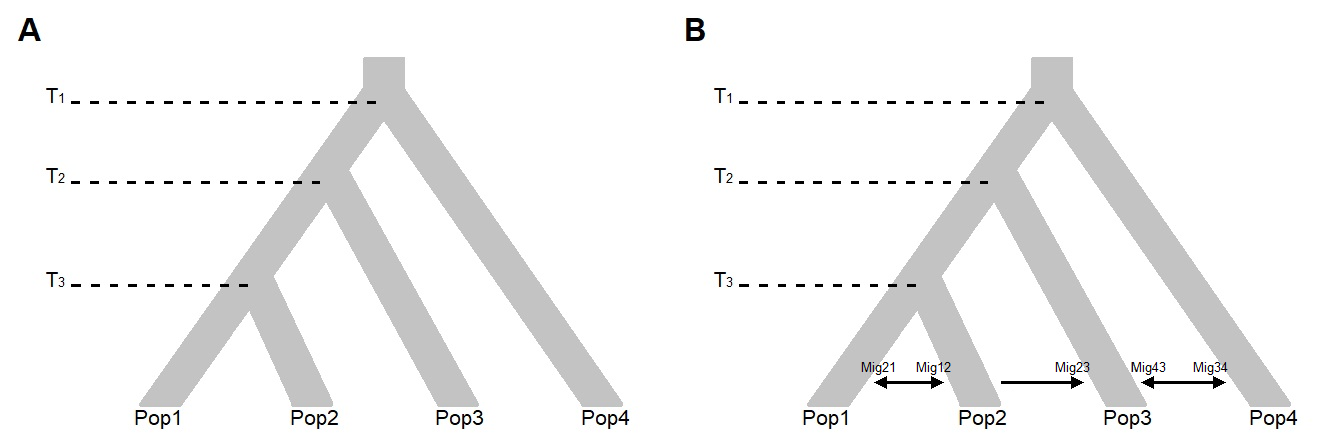

**Figure S6.** Schematic showing the two tested alternative demographic models. In both models, Pop2, Pop3 and Pop4 diverged from Pop1 at the time points T3, T2 and T1 respectively. Panel A shows the null model assuming no secondary contact among populations, in which no migration was allowed among populations. Panel B shows the alternative model with secondary contact among populations, where migration routes were modelled as depicted by the black arrows.
